# BisMapR: a strand-specific, nuclease-based method for genome-wide R-loop detection

**DOI:** 10.1101/2021.01.22.427764

**Authors:** Phillip Wulfridge, Kavitha Sarma

## Abstract

R-loops are three stranded nucleic acid structures with essential roles in many nuclear processes. However, their unchecked accumulation as seen in some neurodevelopmental diseases and cancers and is associated with compromised genome stability. Genome-wide profiling of R-loops in normal cells and their comparison in disease states can help identify precise locations of pathogenic R-loops and advance efforts to attenuate deviant R-loops while preserving biologically important ones. Toward this, we have developed an antibody-independent R-loop detection strategy, BisMapR, that combines nuclease-based R-loop isolation with non-denaturing bisulfite chemistry to produce high-resolution, genome-wide R-loop profiles that retain strand information. Furthermore, BisMapR achieves greater resolution and is faster than existing strand-specific R-loop profiling strategies. We applied BisMapR to reveal discrete R-loop behavior at gene promoters and enhancers. We show that gene promoters exhibiting antisense transcription form R-loops in both directions. and uncover a subset of active enhancers that, despite being bidirectionally transcribed, form R-loops exclusively on one strand. Thus, BisMapR reveals a previously unnoticed feature of active enhancers and provides a tool to systematically examine their mechanisms in gene expression.

## Introduction

R-loops are three-stranded nucleic acid structures that frequently occur during transcription when newly transcribed RNA base pairs with the DNA template strand, forming a DNA:RNA hybrid (Thomas et al., 1976). The non-template strand is then extruded as single-stranded DNA (ssDNA). R-loops play important roles in many nuclear processes including recombination, transcription termination, and DNA repair (Chedin, 2016; Niehrs and Luke, 2020). R-loop formation in mitosis ensures faithful chromosome segregation (Kabeche et al., 2018). While R-loops at some genomic sites clearly have beneficial roles, their aberrant accumulation at others is associated with genomic instability and disease (Crossley et al., 2019; Garcia-Muse and Aguilera, 2019; Perego et al., 2018; Richard and Manley, 2017). The evident role of R-loops in both cellular function and disease makes it critical that reliable methods exist for profiling their formation across the genome. These would allow for detection of changes in R-loop levels between conditions and for the characterization of features associated with their formation that could prove critical towards understanding of disease and development of potential therapies.

Current methods to detect R-loops genome-wide rely on the S9.6 antibody (Dumelie and Jaffrey, 2017; Ginno et al., 2012; Nadel et al., 2015; Wahba et al., 2016) or a catalytically inactive RNase H, both of which recognize DNA:RNA hybrids (Chen et al., 2017; Ginno et al., 2012; Yan et al., 2019). DNA-RNA immunoprecipitation-sequencing (DRIP-seq) is the most frequently used S9.6 based approach for R-loop detection genome-wide (Ginno et al., 2012). In DRIP, genomic DNA is sheared by enzymatic digestion or sonication and regions that contain R-loops are immunoprecipitated using S9.6 antibody. S9.6 based approaches require high input material, have low signal-to-noise ratio and with the exception of BisDRIP-seq (Dumelie and Jaffrey, 2017), have limited resolution. The low signal-to-noise ratio in S9.6 genomewide experiments may be attributed to the antibody specificity issues that are well documented (Hartono et al., 2018; Vanoosthuyse, 2018). RNase H methods that include DNA:RNA in vitro enrichment (DRIVE) (Ginno et al., 2012), R-loop chromatin immunoprecipitation (R-ChIP) (Chen et al., 2017), and MapR (Yan et al., 2019), use the evolutionary specificity of the E.coli RNase H enzyme that recognizes DNA:RNA hybrids to detect R-loops. While similar to DRIP in the initial steps of sample processing, the in vitro enrichment of R-loops in DRIVE using recombinant catalytically inactive RNase H is inefficient. DRIPc (Sanz and Chedin, 2019; Sanz et al., 2016) and R-ChIP are both strand specific techniques with some limitations. DRIPc requires much larger input amounts compared to DRIP and has lower resolution. R-ChIP, a chromatin immunoprecipitation-based strategy, requires the generation of a stable cell line that expresses a catalytic mutant RNase H1 (Chen et al., 2019; Chen et al., 2017; Sanz and Chedin, 2019; Sanz et al., 2016), and while sensitive, may not be amenable for use in all cell types. Therefore, the development of a high-resolution strand-specific R-loop detection strategy that is efficient, sensitive, and amenable to use in all cell types will help in the precise identification of specific regions of the genome that show R-loop anomalies in various diseases.

We recently described MapR (Yan and Sarma, 2020; Yan et al., 2019), a fast and sensitive R-loop detection strategy founded on the principles of CUT&RUN (Skene et al., 2018; Skene and Henikoff, 2017) and the specificity of RNase H for the recognition of DNA:RNA hybrids. In MapR, a catalytically inactive RNase H targets micrococcal nuclease to R-loops to cleave and release them for high throughput sequencing. Because MapR is not enrichment-based, unlike DRIP, DRIVE, and R-ChIP, it has high signal-to-noise ratios resulting in enhanced sensitivity (Yan et al., 2019). We sought to build on the significant advantages of MapR with respect to specificity, sensitivity, and ease of use and transform it into a strand specific R-loop profiling strategy. Here, we present BisMapR, a high resolution, genome-wide methodology that maps strand specific R-loops.

## Results

### BisMapR identifies strand specific R-loops

Strand specificity is a key feature of *bona fide* R-loops. In MapR, cleaved R-loops diffuse out of the nucleus along with mRNAs and other RNAs that are not part of an R-loop. Thus, specific identification of the RNA strand of R-loops as a means of conferring strandedness is challenging with MapR. Instead, we have devised a method to distinguish the template and non-template DNA components of R-loops released by MapR. We leveraged the chemical property of sodium bisulfite to deaminate cytosines (C) to uracils (U) on exposed, single-stranded DNA (ssDNA) only, while double-stranded nucleic acids are protected from conversion (Gough et al., 1986). In BisMapR, R-loops released by MapR are treated with sodium bisulfite under non-denaturing conditions (Fig. 1). This results in the C-to-U conversion of the ssDNA strand of R-loops. Meanwhile, the DNA within the DNA:RNA hybrid is left intact. Next, a second-strand synthesis step replaces the RNA molecule of the DNA:RNA hybrid with a dUTP-containing DNA strand. Adaptors are then ligated to resultant dsDNA. Treatment with uracil DNA glycosylase (UDG) cleaves all dUTP-containing molecules and the unaltered DNA strand is directionally tagged, PCR amplified, and sequenced. The first-mate reads of the resulting paired-end sequencing data (or all reads for single-end runs) correspond to the DNA:RNA hybrid containing strand and are separated into forward- and reverse-strand tracks.

**Figure 1.**
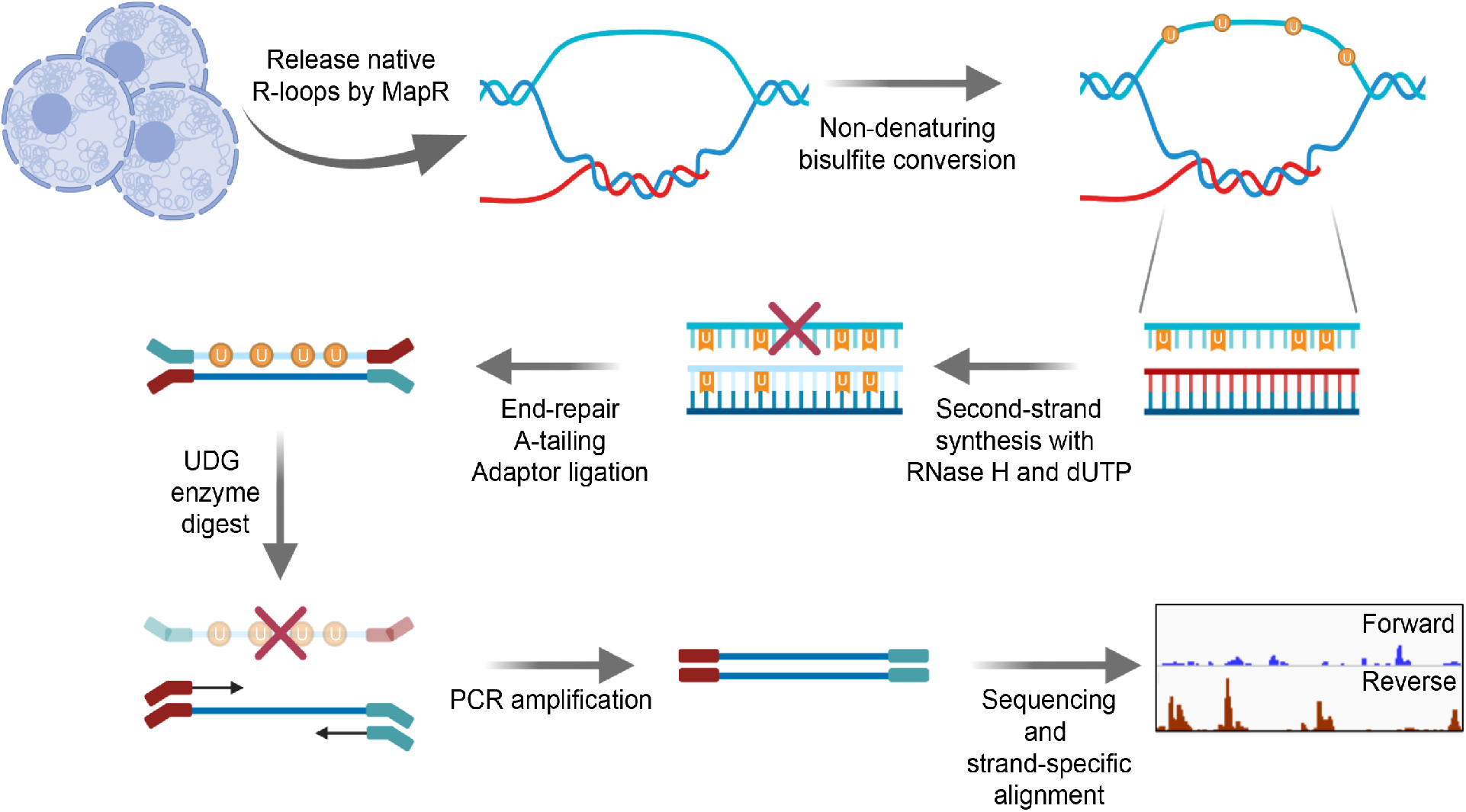
BisMapR, an RNase H based strand-specific native R-loop detection strategy. Schematic of the BisMapR protocol. R-loops are released from cells using MapR and subjected to nondenaturing bisulfite conversion. Bisulfite converted products are directly processed for second-strand synthesis in the presence of RNase H and dUTP. Adaptors are ligated to resultant dsDNA. Treatment with uracil DNA glycosylase (UDG) degrades all uracil containing DNA molecules. Remaining DNA is tagged with paired-end barcoded index primers, amplified, sequenced, and strand-specifically aligned. Created with BioRender.com.

We compared BisMapR and MapR techniques in mouse embryonic stem cells (mESCs). MapR in mESCs specifically identifies R-loops since RNase H catalytic mutant fused to micrococcal nuclease (RH□-MNase) shows high signal at transcription start sites of active genes that are known to form R-loops compared to an MNase-only background control (Supplemental Figs. 1A, 1B). Next, we compared BisMapR to MapR in mESCs to determine if both techniques showed signal enrichment at similar regions within genes. When examining reads from both strands, which we term ‘composite’ signal, BisMapR and MapR produce signal enrichment predominantly around the TSS of actively transcribed genes (Fig. 2A). At the global level also, both datasets showed strong positive correlation for signal enrichment across TSS (Supplemental Fig. 1C). When reads are separated by originating strand, MapR ‘strand-specific’ maps are near-identical to their composite (Fig. 2A), as expected of a non-strand-specific protocol. In contrast, BisMapR reads after strand-specific alignment clearly segregate to either the forward or reverse strands (Fig. 2A). *Vamp1*, a gene that is transcribed in the sense (plus) direction and expected to produce DNA:RNA hybrids on the reverse (template) strand, shows reverse strand specific BisMapR signal. Similarly, *Atg10*, an antisense (minus) gene, shows BisMapR signal on the forward strand. Global analysis of strand-specific BisMapR signal across all TSS also showed decreased correlation with composite BisMapR as well as with strand-specific and composite MapR (Supplemental Fig. 1D).

**Figure 2.**
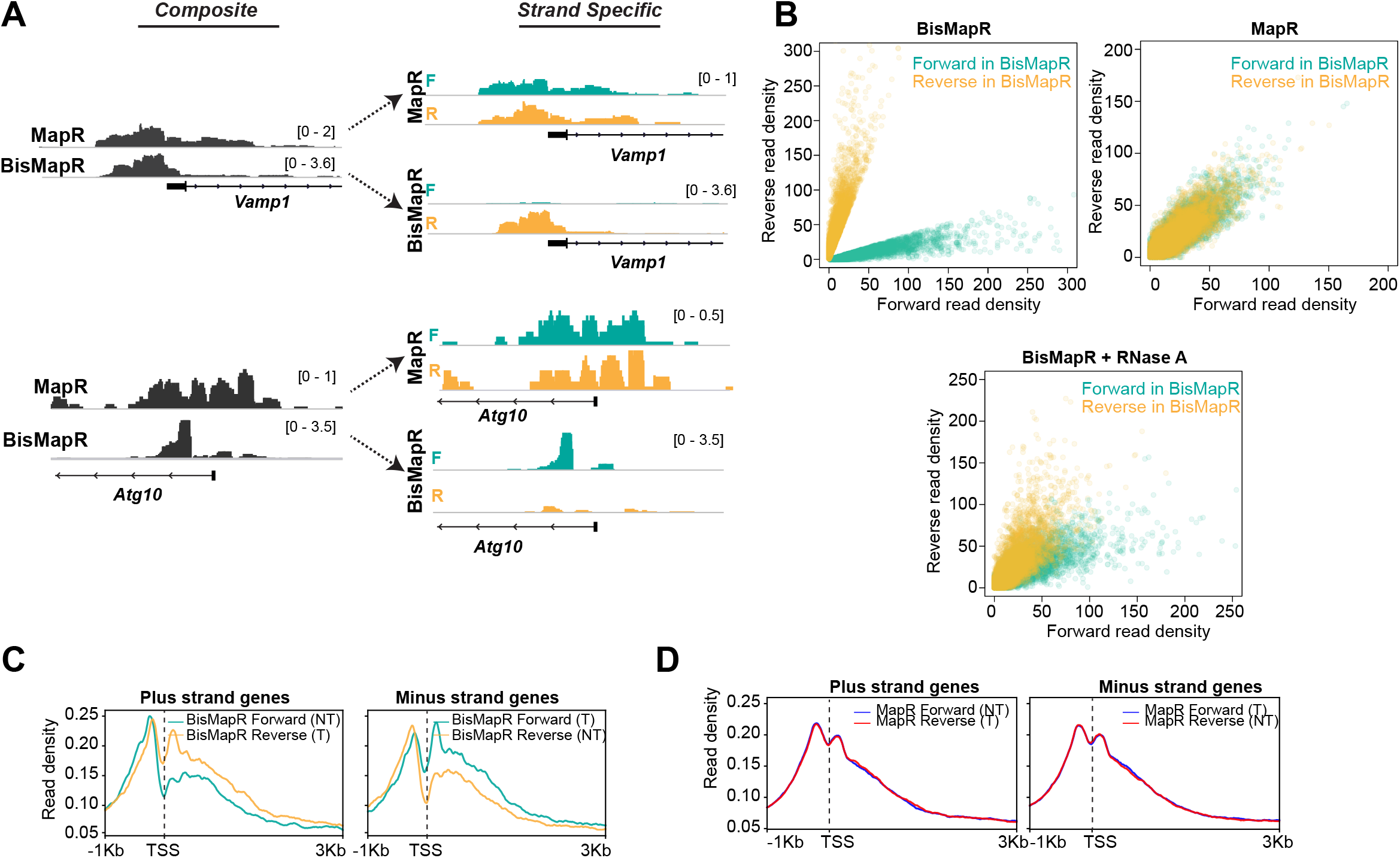
BisMapR confers strand specificity to nuclease-based genome-wide R-loop detection. **A.** Genome browser views of the *Vamp1* and *Atg10* genes showing composite (dark grey) BisMapR and MapR signals (reads per million, RPM) (left) and the same signal when separated into forward (teal) and reverse (orange) strands (right). **B.** Correlation plots of normalized read densities between forward (teal) and reverse (orange) strands in BisMapR, MapR, and BisMapR samples treated with RNase A. TSS regions with a large disparity between forward (teal) and reverse (orange) strand read densities in BisMapR, defined as log2 ratio of at least 1.5 in either direction, are shown. **C.** Metagene plots of BisMapR strand-specific signals at transcription start sites (TSS) of active genes on the plus- and minus-strands in mESCs. Forward strand (teal) and reverse strand (orange) signals are shown. Template (T) and non-template (NT) strands are labeled. **D.** Metagene plots of MapR strand-specific signals at transcription start sites (TSS) of active genes on the plus-strand and minus-strand in mESCs. Template (T) and non-template (NT) strands are labeled.

In MapR, released R-loops are treated with RNase A and the resultant dsDNA that forms as a consequence of degradation of the RNA within the DNA:RNA hybrid is processed for library preparation and sequencing using standard dsDNA library preparation protocols (Yan and Sarma, 2020). However, non-denaturing bisulfite conversion relies on the presence of ssDNA. To assess whether BisMapR requires intact R-loops to efficiently sort signals by strand of origin, we performed bisulfite conversion reactions after treating MapR samples with RNase A and analyzed forward or reverse strand signals across all TSS. BisMapR samples show a clear separation of forward and reverse strands at a large number of TSS in mESCs (Fig. 2B). We observe that MapR, as a non-strand-specific technique, shows little strand separation (Fig. 2B) as seen by the almost complete overlap between forward and reverse strand signals. Treatment of R-loops with RNase A prior to bisulfite processing for BisMapR results in a loss of strand specificity that resembles MapR data, with the forward and reverse strand signals showing a significant overlap (Fig. 2B). The residual strand separation in RNase A treated BisMapR samples is likely due to incomplete RNase A digestion. Thus, BisMapR confers strand specificity on intact R-loops that are released by MapR with minor technical modifications and with a minimal burden on time.

Divergent transcription is a common feature of active promoters in mammals, with over 75% of active genes producing short antisense RNA transcripts and showing paused RNA polymerase II in the antisense direction (Core et al., 2008; Seila et al., 2008). This bidirectional transcription implies that in addition to R-loop formation on the template strand associated with transcription of the gene, R-loops would also form on the opposite strand at these divergent promoters. We used BisMapR to examine R-loops at promoters in mESCs. We confirmed that active genes that likely form R-loops co-transcriptionally show strong R-loop signal around the TSS (Fig. 2C). On the other hand, inactive promoters did not show any measurable BisMapR signal (Supplemental Fig. 1E). As expected from co-transcriptional R-loop formation, BisMapR signal downstream of the TSS is higher on the template strand (T) for both plus and minus-strand genes (Fig. 2C). Additionally, we observe significant signal from the non-template strand (NT) upstream of the TSS, consistent with R-loop function in antisense transcription (Tan-Wong et al., 2019) (Fig. 2C). This clear distinction between template and non-template strand-originating reads cannot be made in MapR, where R-loop signal instead resembles one large peak (Fig. 2D). Thus our data demonstrate that BisMapR produces strand-specific R-loop profiles genome-wide.

### BisMapR can distinguish individual transcriptional units with high resolution

The incorporation of bisulfite treatment that can theoretically react with all single-stranded cytosine residues in the R-loop to mark them for degradation suggests that BisMapR can produce higher resolution R-loop maps compared to MapR. To test this, we analyzed R-loop profiles at active, nonoverlapping head-to-head transcriptional units that are separated by less than a kilobase (Supplemental Fig. 2A). MapR profiles at the 5’ end of *Fbxo18* and *Ankrd16* genes whose TSS are separated by 294 bases show a broad non-strand-specific R-loop enrichment across both TSS (Fig. 3A). In comparison, BisMapR signal is enriched on the forward strand of *Fbxo18* that is transcribed in the antisense direction and on the reverse strand of *Ankrd16* that is transcribed in the sense direction. A similar profile is observed at the *Zfp146* and *Gm5113* genes whose TSS are separated by 125 bases (Supplemental Fig. 2B). Global analysis on all bidirectional transcription units demonstrates that BisMapR captures distinct R-loop signals on opposite template strands that are congruent with the direction of transcription (Fig. 3B). In contrast, MapR signals do not clearly distinguish between pairs of genes (Fig. 3B). Our data suggests that in addition to providing strand-specific information, BisMapR improves on the already high resolution of MapR in profiling R-loops genome-wide.

**Figure 3.**
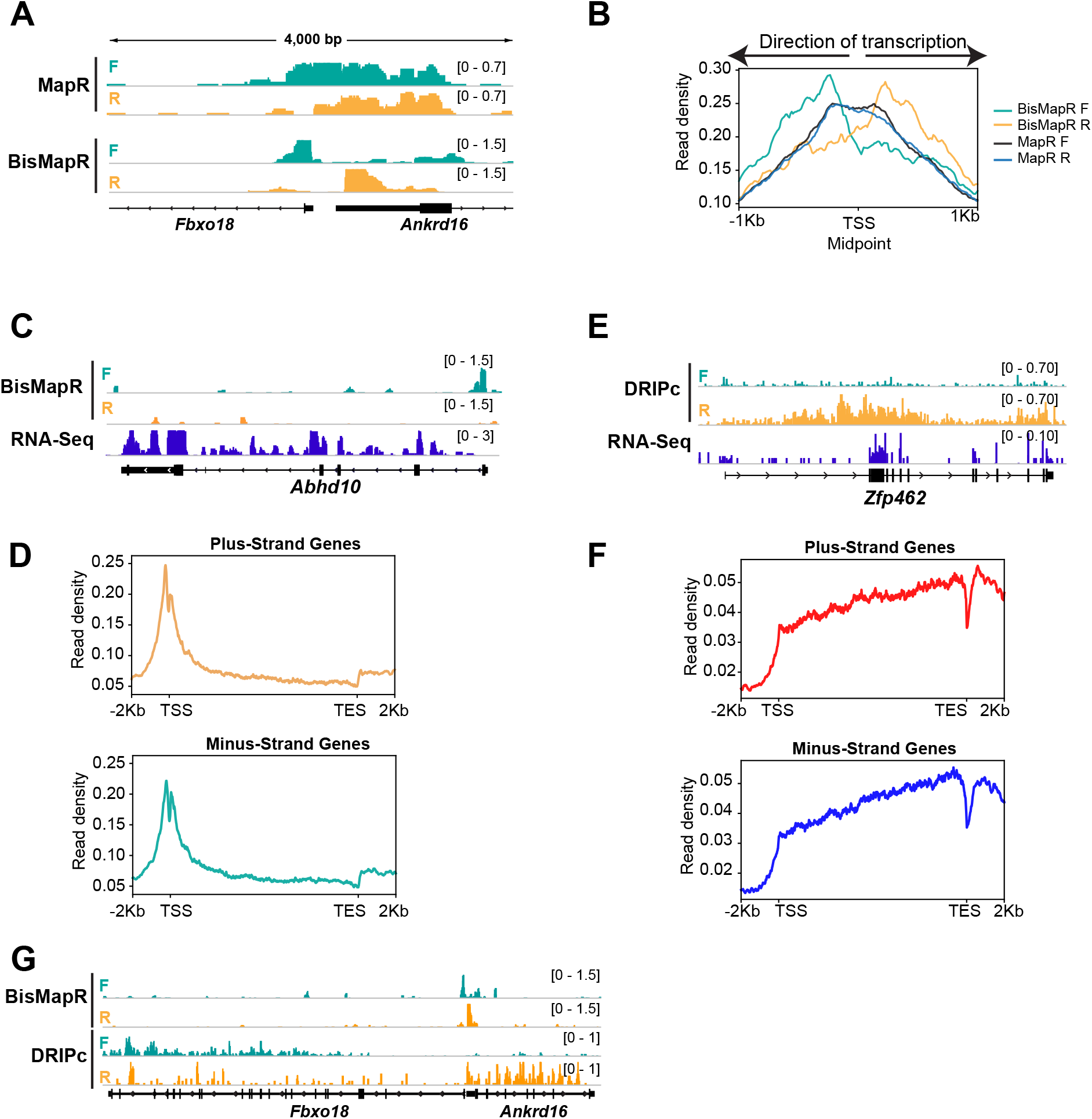
BisMapR distinguishes strand specific R-loops with high resolution at bidirectional gene pairs. **A.** Genome browser view of the bidirectional promoter for *Fbxo18* and *Ankrd16* showing MapR and BisMapR signals (RPM) separated by forward (teal) and reverse (orange) strands. **B.** Metagene plots of strand-specific BisMapR (forward strand (teal) and reverse strand (orange)) and MapR (forward strand (dark blue) and reverse strand (light blue)) signals at bidirectional promoters. TSS Midpoint, the midpoint between the transcription start sites of plus-strand and minus-strand genes. **C.** Genome browser view of the *Abhd10* gene showing BisMapR signal (reads per million, RPM) separated into forward (teal) and reverse (orange) strands. Mouse ESC RNA-seq signal is shown in blue. **D.** Metagene plots of template strand BisMapR signal of plus and minus strand genes expressed in mESCs. **E.** Genome browser view of the *Zfp462* gene showing DRIPc signals separated into forward (teal) and reverse (orange) strands. 3T3 RNA-seq signal is shown in blue. **F.** Metagene plots of template strand DRIPc signal of plus and minus strand genes expressed in 3T3. **G.** Genome browser view of the bidirectional promoter for *Fbxo18* and *Ankrd16* showing BisMapR and DRIPc signals (RPM) separated by forward (teal) and reverse (orange) strands.

Next, we compared the resolution of BisMapR to DRIPc-seq (Sanz and Chedin, 2019; Sanz et al., 2016), an S9.6 antibody-based R-loop enrichment method that is most frequently used to identify strandspecific R-loops. In mESCs, we observe that BisMapR signal is highest at the 5’ end of genes and is reduced across the gene body as is seen in the case of *Abhd10* (Fig. 3C). Metagene analysis across all active genes show that both plus and minus strand genes show 5’ R-loop enrichment (Fig. 3D). This is consistent with several previous reports that observe R-loop enrichment in proximity to the TSS (Chen et al., 2017; Dumelie and Jaffrey, 2017; Ginno et al., 2012; Yan et al., 2019). We used previously published and validated DRIPc data from NIH 3T3 to examine R-loop profiles using this method (Sanz et al., 2016). At *Zfp462*, an expressed gene in NIH 3T3 cells, DRIPc signal is broadly present across the entire gene and does not show a clear enrichment at TSS. Surprisingly, analysis of DRIPc signal profiles across all expressed genes on the plus and minus strands shows that DRIPc-seq shows a general enrichment across transcriptional units on the plus and minus strands starting at the TSS and increasing steadily across the gene body (Fig. 3E).

Our data shows that BisMapR is able to distinguish strand-specific R-loops that form at bidirectional transcription units that are separated by a few hundred bases. To compare the resolution between DRIPc and BisMapR, we visualized R-loops at genes that show head-to-head transcription in both mESCs and NIH3T3 cells. At *Fbxo18* and *Ankrd16*, strand specific BisMapR signal is clearly limited to the TSS of both genes (Fig. 3A, 3G). In contrast, DRIPc signal does not show enrichment at TSS and is instead present across the gene body of both genes (Fig. 3G). Metagene analysis of all active bidirectional transcription units in NIH3T3 shows that DRIPc, while stranded as evidenced by a skew in the forward and reverse strand signals in the appropriate direction, does not delineate two distinct transcriptional units (Supplemental Fig. 2C). Our results indicate that the high resolution of BisMapR, on the order of hundreds of bases, makes it particularly suitable for studying R-loop formation across small-scale features within and between transcriptional units.

### BisMapR reveals unidirectional R-loop formation from KLF7 motifs at a subset of enhancers

Any genomic element with the potential to be transcribed can form R-loops. In addition to genes, enhancers that are transcribed at lower levels and that form short-lived transcripts also form R-loops (Rabani et al., 2014; Schwalb et al., 2016; Yan et al., 2019). To determine differential R-loop formation across enhancers, we examined R-loop signal across active, poised, and primed enhancers in mESCs(Cruz-Molina et al., 2017). Active enhancers are bidirectionally transcribed (Kim et al., 2010; Lai et al., 2015) and show higher global run-on sequencing (GRO-Seq) signals on both the forward and reverse strands as compared to poised and primed enhancers (Supplemental Fig. 3A). Active, poised, and primed enhancers also contain specific chromatin signatures including histone H3 lysine 4 monomethylation (H3K4me1) and histone H3 lysine 27 acetylation (H3K27Ac) at active enhancers, H3K4me1 and H3K27 trimethylation (H3K27me3) at poised enhancers, and H3K4me1 at primed enhancers (Creyghton et al., 2010; Rada-Iglesias et al., 2011; Zentner et al., 2011). We found that active enhancers also exhibit higher MapR and BisMapR signals compared to poised and primed enhancers (Supplemental Fig. 3A, 3B). Interestingly, clustering of all enhancers based on BisMapR forward and reverse strand signals reveals 3 distinct enhancer clusters: two groups with high R-loops that form on either the reverse (group 1) or forward (group 2) strand, and a group with low R-loops (group 3) (Fig. 4A). Both groups of high R-loop enhancers display R-loops on only one strand despite having bidirectional transcription as ascertained by GRO-seq (Fig. 4B). GRO-seq also indicates that group1 and group 2 enhancers are expressed at higher levels than group 3 enhancers. Next, we examined the distribution of active, poised, and primed enhancers across these three groups that we defined based on BisMapR profiles. Group 1 is strongly enriched for active enhancers (70%) and slightly enriched for poised enhancers (0.05%) (Fig. 4C). Similarly, group 2 is also enriched for active (71%) and poised enhancers (0.04%). In contrast, the low R-loop (Fig. 4A) and low transcribed (Fig. 4B) group 3 is enriched for primed enhancers (71%) (Fig. 4C). As expected, genes proximal to group 1 and 2 enhancers, which are enriched for active enhancers, show higher expression levels compared to group 3 enhancer-related genes (Supplemental Fig. 3C). Thus, BisMapR uncovers a subset of enhancers that are transcribed in both directions but exhibit unidirectional strand-specific R-loops.

**Figure 4.**
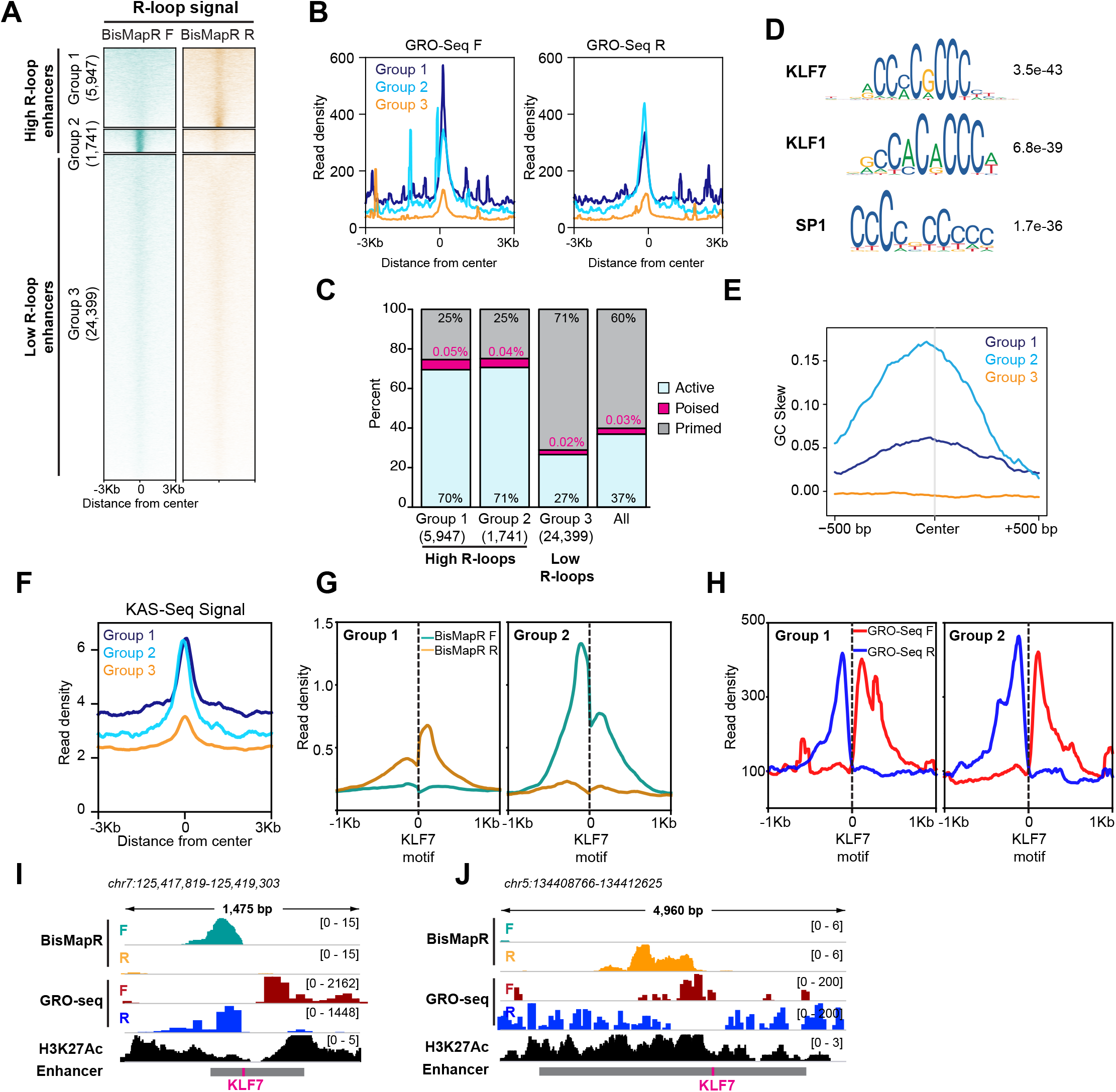
BisMapR reveals strand-specific R-loop formation across a subset of enhancers in mESCs. **A.** Heatmap of strand-specific BisMapR signal across mESC enhancers. Enhancers were divided into 3 groups by unsupervised k-means clustering (k = 3). Enhancer numbers in each group are indicated in parentheses. Signal, reads per million (RPM). **B.** Strand-specific GRO-Seq read densities at Group 1 (dark blue), Group 2 (light blue), and Group 3 (orange) enhancers. **C.** Barplot showing the proportion of active (blue), poised (pink), or primed (grey) enhancers in each group. Enhancer numbers in each group are indicated in parentheses. Distribution of all enhancer types is shown for comparison. **D.** Sequence motifs of KLF7, KLF1, and SP1 transcription factors that are centrally enriched across Group 1 and Group 2 enhancers compared to Group 3 enhancers. E-values for each sequence are shown. **E.** GC skew of non-template strand across Group 1 (dark blue), Group 2 (light blue), and Group 3 (orange) enhancers. GC skew was calculated as (G-C)/(G+C) in 100-bp sliding windows with 10-bp step size. **F.** KAS-Seq read density across Group 1, 2, and 3 enhancers. **G.** Strand-specific BisMapR signal (RPM) profiles centered around Klf7 motifs identified in Group 1 and Group 2 enhancers. **H.** Strand-specific GRO-seq signal profiles centered around Klf7 motifs identified in Group 1 and Group 2 enhancers. **I, J.** Genome browser view of high-R-loop enhancers showing strand-specific BisMapR (F, teal; R, orange) and GRO-seq (F, red; R, blue) signals and H3K27Ac ChIP-seq (black). Enhancer region (grey bar) with location of KLF7 motif (pink) is shown.

Next, we sought to identify distinguishing features between high and low R-loop enhancers. We used CentriMo to identify sequences that are centrally enriched in the two high R-loop enhancer groups (Groups 1 and 2) compared to the low R-loop group (Group 3). Our analysis uncovered a significant enrichment for KLF7, KLF1, and SP1 transcription factor motifs in the high R-loop enhancer groups (Fig. 4D). Since all three motifs appear to have a high GC content, we performed a GC skew analysis to determine if the non-template strand that forms the ssDNA component of R-loops is enriched for guanines (G). We found that the non-template strands in groups 1 and 2 have high G content, while group 3 shows no skew (Fig. 4E). It is possible that the abundance of guanines on the ssDNA can promote the formation of G-quadruplex (GQ) structures that may aid in stabilization of these enhancer R-loops. A recently developed technique, kethoxal-assisted single-stranded DNA sequencing (KAS-seq), identifies single stranded regions of the genome with high G content (Wu et al., 2020). In KAS-seq, an azide-tagged kethoxal (Weng et al., 2020) that reacts with unpaired guanine residues and that can be modified with a biotin is used to mark and enrich for G-rich regions of the genome that are single stranded. We found that high R-loop enhancers as determined by our analysis (Groups 1 and 2) exhibit high KAS-seq signal, consistent with our observation of GC skew and the likelihood of containing ssDNA, a component of R-loops (Fig. 4F). To further determine the degree of concordance between KAS-Seq and BisMapR, we performed clustering of all enhancers based on KAS-seq signal, identifying 5,069 enhancers with high KAS-seq signal and 27,811 with low KAS-seq signal. 2,226 enhancers had both high KAS-seq and high R-loop signals, representing a 1.83-fold enrichment. In contrast, enhancers with high R-loops were under-enriched (1.19-fold) in the enhancer population with low KAS-seq signals. Therefore, BisMapR and KAS-seq present two complementary methods to identify enhancers that contain ssDNA components.

Motif analysis at high R-loop enhancers identified an enrichment for KLF7, KLF1, and SP1 binding sites (Fig. 4D). KLF7, KLF1, and SP1 are pioneer transcription factors that have the ability to bind DNA and open condensed chromatin. Interestingly, analysis of transcription factor binding across DNase I hypersensitive sites showed that KLF7 motif is associated with asymmetrically open chromatin, suggesting a directional pioneer factor activity (Sherwood et al., 2014). To examine whether R-loops are skewed at enhancers that contain KLF7 binding sites, we re-centered enhancer regions in groups 1 and 2 around their KLF7 motifs and examined BisMapR signal (Fig. 4G). Interestingly, in both these groups strandspecific R-loops form unidirectionally from KLF motifs (Fig. 4H, 4I, 4J, Supplemental Fig. 4D). Presence of H3K27 acetylation (Fig. 4I, 4J) and GRO-seq signals (Fig. 4H, 4I, 4J) around the KLF7 centered enhancers show clear bidirectional transcription. Our data suggests that KLF7 binding may contribute to the unidirectional formation or stabilization of R-loops at some active enhancers that show high levels of transcription.

## Discussion

Genome-wide R-loop mapping relies on two distinct approaches, the S9.6 antibody and a catalytically inactive RNase H, that both recognize DNA:RNA hybrids. These two approaches recognize some common R-loops, as well as other unique subsets that likely appear as a consequence of the distinct sequence preferences of S9.6 and RNase H (Konig et al., 2017). Therefore, orthogonal approaches that are sensitive, efficient, and that retain strand specificity will allow for comprehensive interrogation of R-loop dynamics in various cellular process and their dysfunction in disease. Here, we have described BisMapR, a fast RNase H based method that efficiently captures strand-specific R-loops at high resolution genome-wide.

Existing strand-specific R-loop mapping strategies have a few drawbacks that include low resolution and high input material (Sanz and Chedin, 2019; Sanz et al., 2016), involved sample processing (Dumelie and Jaffrey, 2017), and lengthy experiment times (Chen et al., 2019; Chen et al., 2017). Our comparison of BisMapR with DRIPc, an S9.6 based approach, shows that BisMapR produces sharper regions of enrichment suggesting higher resolution (Fig. 3). The high resolution of BisMapR is especially evident at head-to-head transcribed genes TSS that are only a few hundred bases apart and that show R-loop formation on opposite strands. As noted before (Sanz and Chedin, 2019), a reason for the broader signal seen in DRIPc can be because of immunoprecipitation of the regions of the RNA that are not part of the R-loop. While non-denaturing bisulfite treatment can potentially correct this drawback, DRIPc still requires a significantly higher amount of starting material compared to DRIP and MapR. Bisulfite conversion of DRIP samples *after* immunoprecipitation may help reduce input requirement and achieve higher resolution and provide an S9.6 based approach that is on par with the resolution and sensitivity of BisMapR.

Using BisMapR, we uncovered a subset of enhancers with high R-loops and that have a high GC skew. R-loop formation is associated with the potential to form G quadruplexes on the non-template strand. Two recent studies using single molecule approaches have provided insight into how G quadruplexes and R-loop formation regulate gene expression. While R-loop formation precedes GQ formation, stable GQs in the non-template strand provide a positive feedback to promote R-loops during transcription (Lim and Hohng, 2020). Transcription efficiency is increased as a result of successive rounds of R-loop formation (Lee et al., 2020). Taken together with our finding of high R-loops at some enhancers, it is possible that R-loop formation and GQ stabilization leads to the maintenance of nucleosome free regions and help in sustained enhancer activation.

In summary, BisMapR is a fast, sensitive, and strand-specific R-loop detection strategy that reveals strand specific R-loops at enhancers that are also enriched for KLF7 pioneer factor binding motifs. Our study provides a tool to further dissect how directional chromatin accessibility conferred by a subset of pioneer factors contributes to R-loop stabilization at enhancers and their combined significance to gene expression.

## Acknowledgements

We thank A. Gardini and R. Bonasio for helpful discussions, and Q. Yan for assistance with MapR experiments. This work was supported by National Institutes of Health Grants DP2-NS105576 (to K.S.) and T32CA009171 (to P.W.).

## Methods

### Cell culture

E14 mouse embryonic stem cells (mESCs) were cultured on 0.1% gelatin coated plates in media containing DMEM, 15% fetal bovine serum (Gibco), 1 x MEM non-essential amino acids, 1X GlutaMAX (Gibco 35050), 25mM HEPES, 100U/ml Pen-Strep, 55μM 2-mercaptoethanol, 3μM glycogen synthase kinase (GSK) inhibitor (Millipore 361559), 1μM MEK1/2 inhibitor (Millipore 444966), and LIF (Sigma, ESGRO).

### BisMapR

MapR was performed as described (Yan and Sarma, 2020; Yan et al., 2019) on 5×10^6^ cells for BisMapR and control MapR samples (n = 2 replicates per method), with the exception that RNase A was omitted from the stop buffer of the BisMapR sample. Following DNA extraction, the BisMapR samples were bisulfite converted using reagents from the EZ DNA Methylation-Gold Kit (Zymo D5005). 10 μL of sample was added to 10 μL of dH_2_O and 130 μL of CT Conversion Reagent, then incubated for 3 hours at room temperature (25°C) to preserve double-stranded nucleic acids under non-denaturing conditions. DNA desulfonation and column purification was performed according to manufacturer’s instructions and eluted into 20 μL of M-Elution buffer. The elution product was directly used for second-strand synthesis using reagents from the NEBNext Ultra II Directional RNA Library Prep Kit (NEB E7760). 8 μL of Second Strand Synthesis Reaction Buffer, 4 μL Second Strand Synthesis Enzyme Mix, and 48 μL dH_2_O was added to elution product, and the reaction was incubated for 1 hour at 16°C in a thermocycler. Double-stranded DNA was purified from the second-strand reaction using 1.8x volume of AMPure XP SPRI beads (Beckman Coulter). As a negative control, BisMapR bisulfite conversion and second-strand synthesis steps were also performed on a MapR sample in which RNase A was present in the stop buffer (+RNase A).

### RNA-Seq

RNA samples were extracted from mESCs (n = 3 biological replicates) using Trizol reagent (Invitrogen) and subjected to DNase digestion with Turbo DNase (Ambion AM2238). RNA samples were then rRNA-depleted using FastSelect-rRNA HMR (Qiagen) and converted to cDNA using Ultra II Directional RNA Library Prep Kit (NEB E7760).

### Library preparation and sequencing

DNA samples were end-repaired using End-Repair Mix (Enzymatics), A-tailed using Klenow exonuclease minus (Enzymatics), purified with MinElute columns (Qiagen), and ligated to Illumina adaptors (NEB E7600) with T4 DNA ligase (Enzymatics). Size selection for fragments >150 bp was performed with AMPure XP (Beckman Coulter). Libraries were PCR amplified with dual index barcode primers for Illumina sequencing (NEB E7600) using Q5 DNA polymerase (NEB M0491) and purified with MinElute. Uracil DNA glycosylase (Enzymatics) was added to the PCR amplification mix to degrade dUTP-containing molecules and remove adaptor hairpins. Sequencing was performed on a NextSeq 500 instrument (Illumina) with 38×2 paired-end cycles.

### Data processing

BisMapR, MapR, and DRIPc-seq reads were mapped to the mouse genome (mm10) with Bowtie2 version 2.2.9 (Langmead and Salzberg, 2012) using default parameters and paired-end (BisMapR and MapR) or single-end (DRIPc-seq) settings as appropriate. Mouse ESC and NIH3T3 RNA-Seq reads were mapped to mm10 with STAR version 2.7.3 (Dobin et al., 2013) and RSEM (Li and Dewey, 2011) version 1.3.3 was used to obtain estimated counts. A gene was considered expressed if it had at least 1 count per million (CPM) in all RNA-Seq samples. A bidirectional promoter was defined as a region 1 kb or smaller containing two transcription start sites for genes in opposite directions, i.e. sense and anti-sense, and with both genes expressed in mESC or NIH3T3. To generate strand-specific datasets for BisMapR, MapR, RNA-Seq, and DRIPc-seq, reads were separated based on the strand to which the first mate aligned. Specifically, reads with SAM flags 16, 83, or 163 were placed into the forward-strand dataset, while reads with SAM flags 0, 99, or 147 were placed into the reverse-strand dataset. The first-mate strand represents the template strand for BisMapR, MapR, and DRIPc-seq. RPM normalization for strand-specific datasets was calculated based on the combined number of reads assigned to either the forward or reverse strand. BigWig tracks were generated using the bamCoverage function in deepTools 3.4.1 (Ramirez et al., 2016) with options-- binSize 5 and --blackListFileName to remove a known set of ENCODE blacklist regions (Amemiya et al., 2019). The --extendReads option was used for paired-end datasets. Signal plots and heatmaps were generated using the computeMatrix, plotProfile, and plotHeatmap functions in deepTools. Motif enrichment analysis was performed with CentriMo using 500-bp sequences centered around each enhancer.

### Published data

We downloaded FASTQ files for NIH3T3 DRIPc-seq (SRR3322169) and NIH3T3 RNA-seq (SRR6126847), BigWig tracks for KAS-seq (Wu et al., 2020) (GSE139420), GRO-seq (Tastemel et al., 2017) (GSE99760) and H3K27Ac (ENCODE ENCFF163HEV), and enhancer locations from (Cruz-Molina et al., 2017).

### Data availability

Sequencing data generated for this study have been deposited in the NCBI GEO as GSE160578. Data will remain private during peer review and released upon publication.

## Supplemental figure legends

**Supplemental Figure 1.**
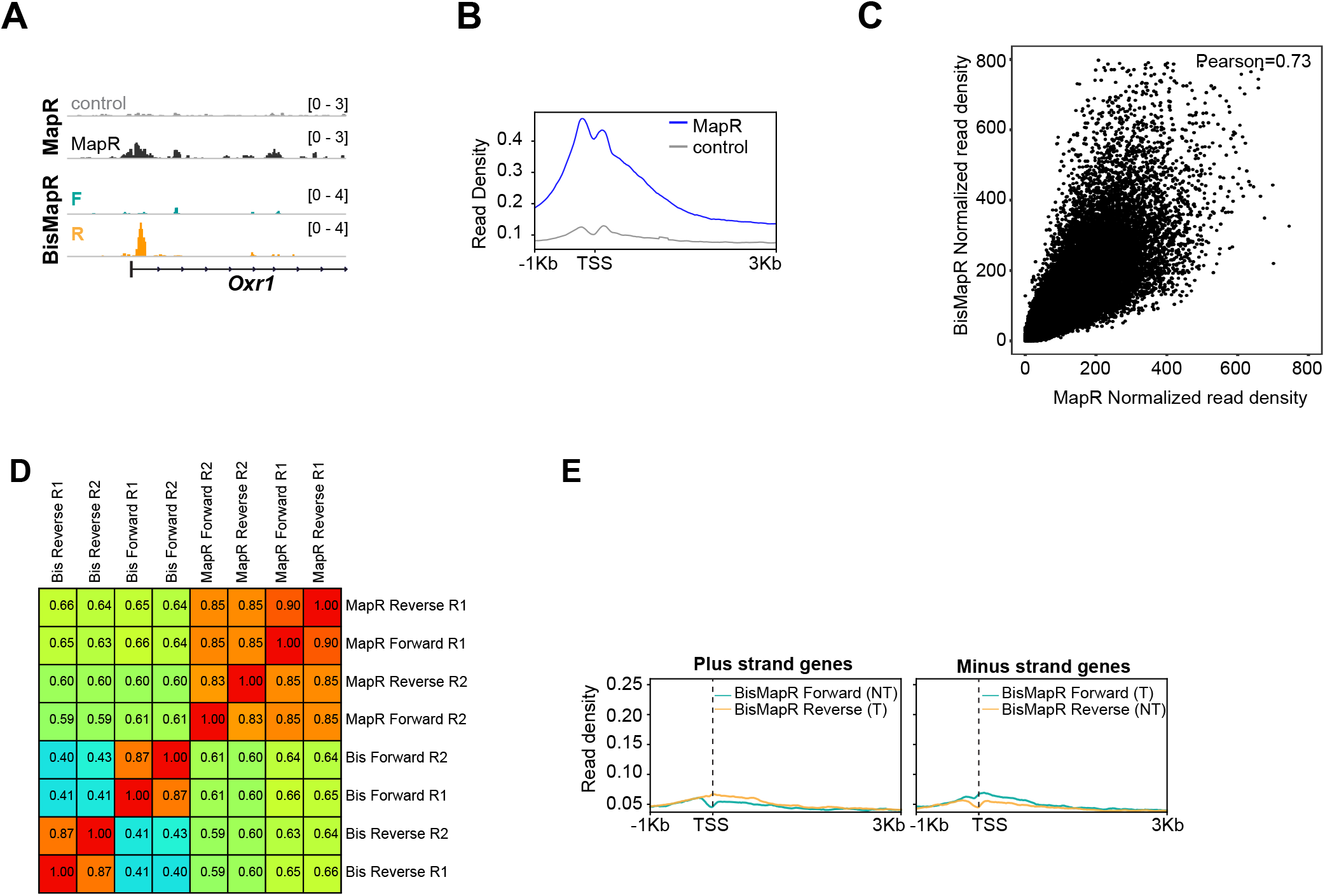
BisMapR generates strand-specific transcription dependent R-loop sequencing data. **A.** Genome browser view of the *Oxr1* gene showing GST-MNase (control) and GST-RH□-MNase (MapR) composite signals and BisMapR strand specific signals. Forward, teal. Reverse, orange. **B.** Metagene plot of control and MapR signals at all active TSS in mESCs. **C.** Correlation plot showing normalized read density between MapR composite and BisMapR composite samples. Pearson correlation, 0.73. For all correlation analysis plots, read density was calculated for 1kb flanking regions across all mouse TSS. **D.** Heatmap showing pairwise Pearson correlation scores of composite, forward-strand, and reverse-strand BisMapR and MapR read alignments. **E.** Metagene plots of BisMapR strand-specific signals at transcription start sites (TSS) of inactive genes on the plus- and minus-strands in mESCs. Forward strand (teal) and reverse strand (orange) signals are shown. Template (T) and non-template (NT) strands are labeled.

**Supplemental Figure 2.**
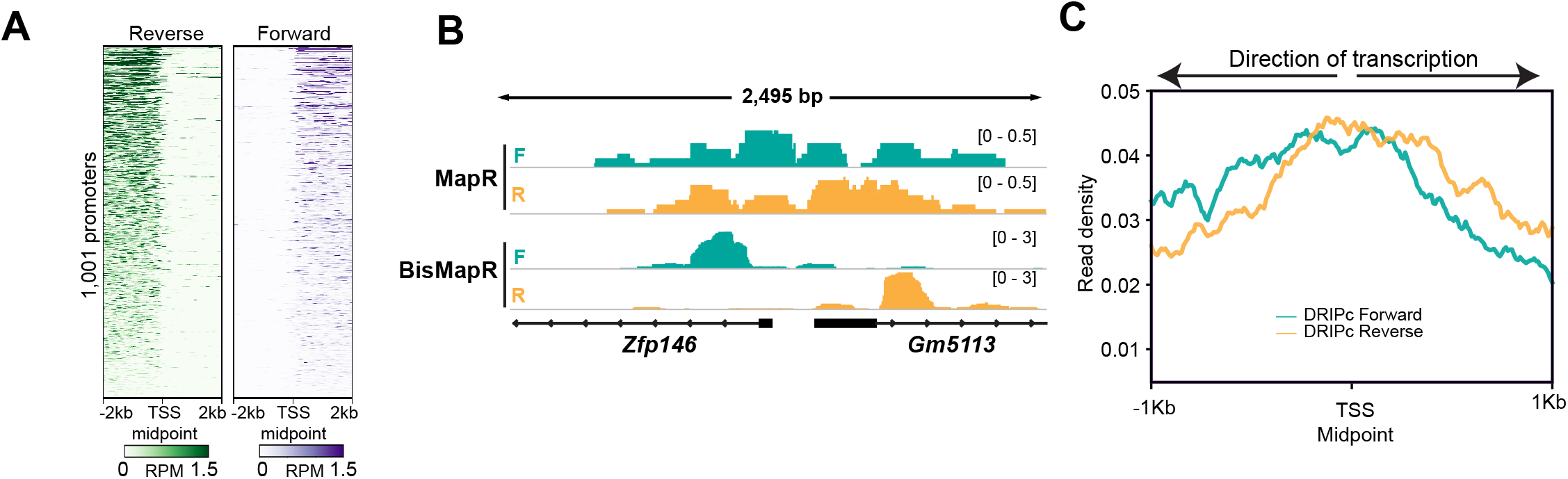
BisMapR reveals strand specificity of R-loops at high resolution at genic regions genome-wide. **A.** Heatmap of RNA-Seq strand-specific reads at bidirectional promoters. All rows are oriented in the plus-strand direction. Midpoint between the transcription start sites of plus-strand and minus-strand genes is shown. Signal, reads per million (RPM). **B.** Genome browser view of the bidirectional promoter for *Zfp146* and *Gm5113* showing strand-specific MapR and BisMapR signal (RPM). **C.** Metagene plots of strand-specific DRIPc-seq (Forward strand (teal) and reverse strand (orange)) signal at 3T3 bidirectional promoters. TSS Midpoint, the midpoint between the transcription start sites of plus-strand and minus-strand genes.

**Supplemental Figure 3:**
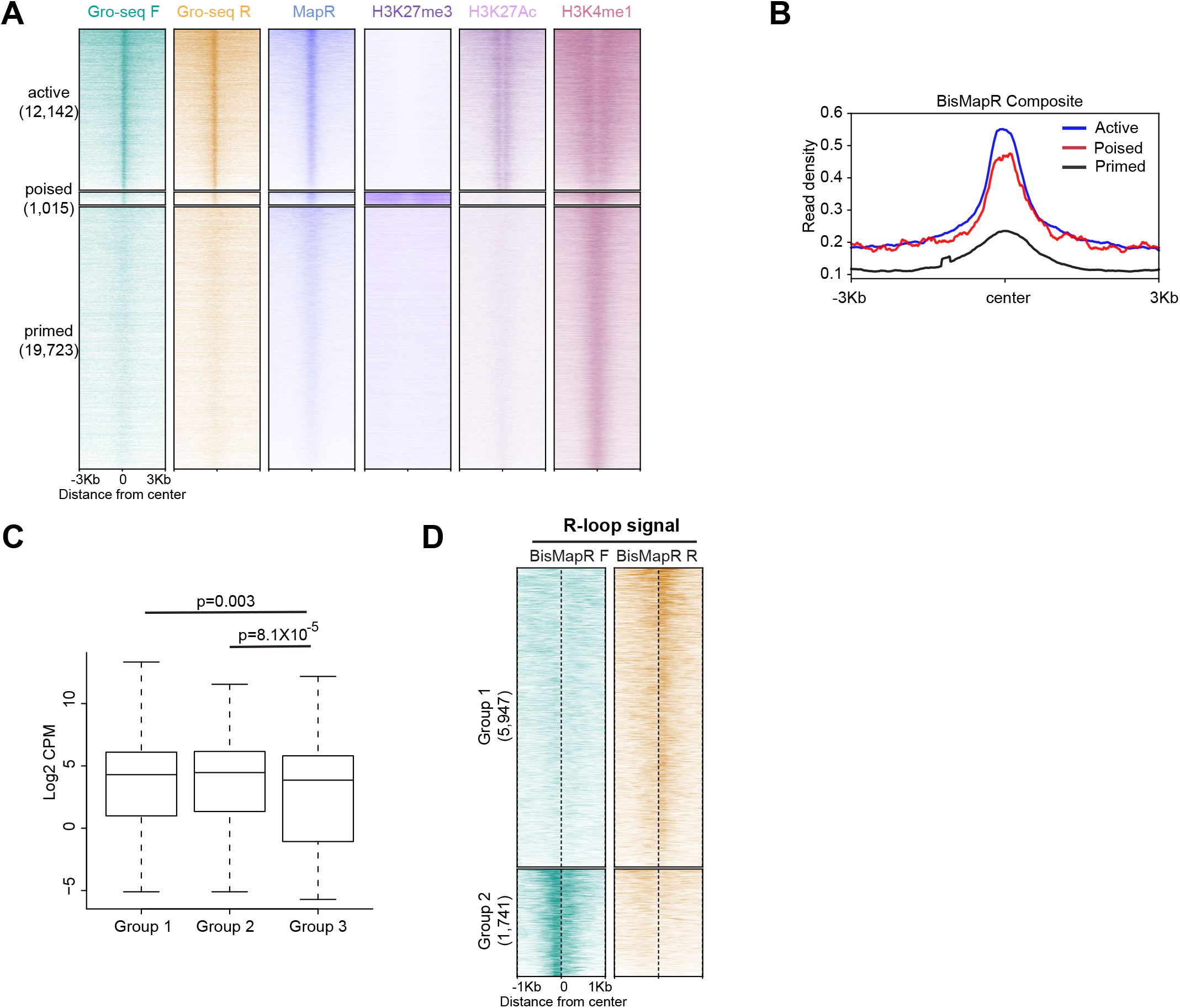
BisMapR identifies strand specific R-loops at enhancers. **A.** Heatmap of strand-specific GRO-Seq read density, MapR, H3K27me3, H3K27Ac, and H3K4me1 at active, poised, and primed mESC enhancers. Enhancers were divided into active, poised, and primed. Enhancer numbers in each group are indicated in parentheses. **B.** BisMapR composite signal profiles at active (blue), poised (red), and primed (black) enhancers. **C.** Boxplot showing mESC expression levels of genes associated with enhancers in Groups 1, 2, or 3. Welch’s t-test p-values are shown. **D.** Heatmap of strand-specific BisMapR signal centered at KLF7 binding motifs in Group 1 and Group 2 enhancers. Enhancer numbers in each group are indicated in parentheses. Signal, reads per million (RPM).

## Notes

### Competing Interest Statement

The authors have declared no competing interest.

